# Longevity and rejuvenation effects of cell reprogramming are decoupled from loss of somatic identity

**DOI:** 10.1101/2022.12.12.520058

**Authors:** Dmitrii Kriukov, Ekaterina E. Khrameeva, Vadim N. Gladyshev, Sergey E. Dmitriev, Alexander Tyshkovskiy

**Author notes:** Correspondence should be addressed: A.T., S.E.D.

## Abstract

Partial somatic cell reprogramming has been touted as a promising rejuvenation strategy. However, its association with mechanisms of aging and longevity at the molecular level remains unclear. We identified a robust transcriptomic signature of reprogramming in mouse and human cells that revealed co-regulation of genes associated with reprogramming and response to lifespan-extending interventions, including those related to DNA repair and inflammation. We found that age-related gene expression changes were reversed during reprogramming, as confirmed by transcriptomic aging clocks. The longevity and rejuvenation effects induced by reprogramming in the transcriptome were mainly independent of pluripotency gain. Decoupling of these processes allowed predicting interventions mimicking reprogramming-induced rejuvenation (RIR) without affecting somatic cell identity, including an anti-inflammatory compound osthol, *ATG5* overexpression, and *C6ORF223* knockout. Overall, we revealed specific molecular mechanisms associated with RIR at the gene expression level and developed tools for discovering interventions that support the rejuvenation effect of reprogramming without posing the risk of neoplasia.

## Introduction

Aging is associated with the buildup of molecular damage and a gradual loss of function, culminating in chronic age-related diseases and ultimately death (*1*). Searching for safe and efficient interventions that can slow down or partially reverse the aging process is a major challenge in the aging field (*2–6*). In this regard, reprogramming of somatic cells into induced pluripotent stem cells (iPSCs) has been proposed as a candidate longevity intervention due to its potential to rejuvenate cells in a targeted way (*7, 8*).

Pluripotency can be achieved *in vitro* by the ectopic expression of four transcription factors: OCT4, SOX2, KLF4, and MYC, known as OSKM or Yamanaka factors (YFs). It was demonstrated that OSKM support the generation of murine iPSCs (*9*) using retroviral transduction as a delivery system and mouse embryonic fibroblasts (MEF) as the initial cell culture. Although this original experiment was inefficient in terms of the percentage of cells that terminally achieved the pluripotent state (*<*0.1%), more advanced *in vitro* approaches resulted in a greatly improved efficiency, e.g. by down-regulation of methyl CpG-binding domain 3 (MBD3) levels (*10*). In parallel, other approaches have been developed to induce pluripotency. In particular, the expression of seven other transcription factors (7F: *Jdp2-Jhdm1b-Mkk6-Glis1-Nanog-Essrb-Sall4*) resulted in high efficiency of reprogramming (*11*). Therefore, it appears that the reprogramming process can be attained by using different cell culture conditions, transcription factors, and small molecules (*12*).

*In vivo* cell reprogramming could be accomplished by using transgenic mice with doxycycline-inducible OSKM (*13, 14*). However, continuous expression of OSKM factors in mice leads to severe forms of teratoma. Partial reprogramming protocols can overcome this problem. Some of these techniques rely on the incomplete set of reprogramming factors, e.g., OSK reprogramming (*15*), while others are based on a transient or temporarily controlled expression of OSKM factors (*16–18*). The problem of oncogenesis during *in vivo* reprogramming is associated with the loss of somatic cell identity in pluripotent cells. Thus, it is crucial to avoid the reset of the somatic epigenetic program in order to make this technique applicable in clinical practice. Recent *in vitro* experiments (*19, 20*) show that the decrease in epigenetic age of reprogrammed cells measured by epigenetic aging clocks (*21*) occurs mostly prior to the stabilization phase when the pluripotent state is established. However, even a short-term use of OSKM factors has been shown to produce the detectable subpopulation of cells where late-stage pluripotent genes are expressed (*22*). Therefore, independent interventions that would support Reprogramming-Induced Rejuvenation (RIR) without inducing pluripotency may be of a high clinical value.

Since the first reprogramming experiment conducted in 2006, a massive amount of high-throughput molecular data have accumulated, shedding light on the details of gene regulatory pathways and their dynamics during reprogramming. These data comprise transcriptome, methylome, chromatin conformation, chromatin accessibility, and other omics datasets, including single-cell transcriptomes (*22, 23*). The resulting data on reprogramming allowed the construction of various mathematical models (*24–26*) describing some aspects of reprogramming. However, models that offer specific molecular mechanisms responsible for RIR have been lacking.

Here, we describe transcriptomic changes that occur in cells during reprogramming and their association with mechanisms of aging and longevity. We conducted a comprehensive meta-analysis of time-course gene expression datasets of mouse and human cells during multi-factor reprogramming and identified robust transcriptomic signatures associated with this process. By integrating them with the signatures of aging and lifespan-extending interventions, we revealed genes and functional processes specifically associated with the rejuvenation and longevity effects of reprogramming. Using multi-tissue transcriptomic aging clocks developed for humans and mice, we further observed a significant reduction of the transcriptomic age (tAge) for both human and mouse cells in response to OSKM, OSK and 7F reprogramming. Remarkably, most genes responsible for the rejuvenation and longevity effects of reprogramming were not involved in the loss of somatic identity and gain of pluripotency, suggesting that these processes can be separated. This allowed us to identify specific gene expression signatures of RIR and use them to discover candidate chemical and genetic interventions that may induce reprogramming-associated rejuvenation effects without affecting somatic cell identity.

## Results

### Reprogramming gene expression signature captures dynamics towards pluripotency

We gathered 41 gene expression datasets of time-course cell reprogramming from 14 studies, including 29 datasets for mouse cells and 12 datasets for human cells (Suppl. Table S1, Suppl. Fig. S1). Each dataset represented a continuous cell reprogramming experiment conducted on a specific cell line with a particular treatment, including OSKM, OSK or 7F. Most murine datasets were MEF-iPSC reprogramming, whereas human studies used different types of cells. To identify genes, whose expression was robustly changed during the reprogramming process across the datasets, we utilized mixed-effect linear models previously used to discover transcriptomic signatures of lifespan-extending interventions and aging (see Methods, Fig. 1A, (*27*)).

**Fig. 1.**
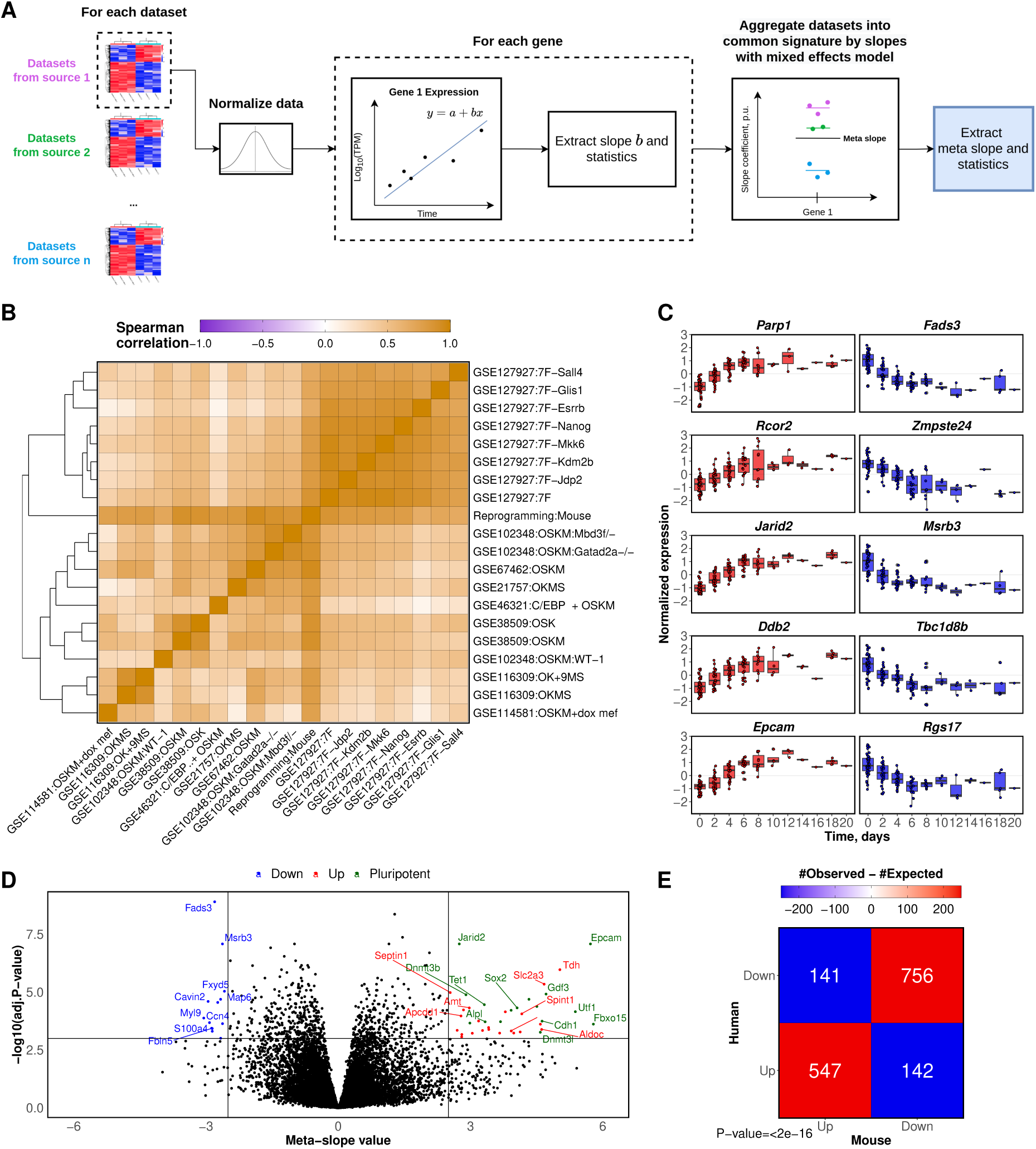
Construction and validation of the reprogramming signature. **(A)** Schematic illustration of the signature construction workflow. **(B)** Clustering analysis of individual mouse reprogramming datasets and the aggregated signature. GSE IDs of the datasets are accompanied by the description of reprogramming factors applied in corresponding experiments (for details, see Table S1). Cells are colored based on Spearman’s correlation coefficient. **(C)** Expression trajectory of top five upregulated and downregulated genes with the lowest BH-adjusted p-value according to the mouse reprogramming signature. Upregulated and downregulated genes are shown in red and blue, respectively. **(D)** Volcano plot of meta-slope values extracted from the signature and corresponding BH-adjusted p-values. Each dot represents a single gene. Pluripotency markers are highlighted in green. Significantly upregulated and downregulated genes are shown in red and blue, respectively. The horizontal dashed line represents the significance cut-off (BH adjusted p-value < 0.001), while vertical lines represent meta-slope cut-offs (|logFC| > 2.5). **(E)** The overlap of significantly upregulated and downregulated genes between murine and human signatures. Only uniquely mapped orthologs according to Ensembl were considered for analysis. Numbers within cells demonstrate the observed numbers of overlapping orthologous genes, while color represents the difference between observed and expected number of genes in the corresponding cell. The p-value is calculated using Fisher’s exact test.

Using this approach, we identified mouse- and human-specific gene expression signatures of reprogramming, as well as common reprogramming signatures conserved across species. In total, 3087, 7531, and 4807 genes changed their expression in the cells from mice, humans, and both species, respectively, during reprogramming (BH-adjusted p-value < 0.05). A higher number of significant genes for humans may be related to batch effects, since all corresponding time-course datasets have been created by the same research group. Therefore, in this study we mostly focus on the mouse signature, as it offers more reproducible and robust biomarkers of cellular reprogramming.

To assess the quality of signatures, we checked if they recapitulate gene expression changes in individual datasets used for their construction (Fig. 1B, Suppl. Fig. S2A). Both murine and human signatures demonstrated a significant positive Spearman correlation with each utilized dataset (rho > 0.66). Clusters were generally formed by datasets from the same source (i.e., the same GSE ID) and based on the same type of treatment. Thus, classical Yamanaka factors (YF), including OSKM and OSK treatments, clustered together and were separate from the 7F intervention (*11*). In addition, human datasets clustered mainly by tissue type (Suppl. Fig. S2A). Overall, the correlation analysis suggested that the constructed reprogramming signature captured consistent gene expression changes observed in multiple independent experiments.

To investigate the expression dynamics of top genes associated with reprogramming, we visualized normalized expression of 5 up- and 5 downregulated genes with the lowest p-values (Fig. 1C, Fig. S2B). We observed saturation of gene expression dynamics after 10 days of reprogramming in mice, while in human cells top genes demonstrated sigmoid-like dynamics with the saturation point at 20th day. Interestingly, several top up- and downregulated genes captured by our signatures were previously shown to be associated with aging and included in the GenAge database (*28*). Thus, *Parp1* is known as an antagonistic pleiotropic gene (*29*) regulating genome maintenance and inflammation processes. At the same time, in cooperation with *Sox2* it can function as an alternative splicing regulator during reprogramming (*30*). Another example is the downregulated *Zmpste24* gene, whose deficiency in mice results in nuclear architecture abnormalities and signs of accelerated aging (*31*).

For additional validation of our signatures, we examined the distribution of pluripotency-associated genes (see Table S2 for a full list of pluripotency genes) among those significantly perturbed during reprogramming (Fig. 1D, Fig. S2C and S2D; see Table S3 for the list of top genes in the signature). Notably, significantly upregulated genes (BH-adjusted p-value < 0.001) were enriched by the markers of pluripotency (Fisher exact test p-value *<* 1*^−^*^10^). In particular, *Zic3, Gdf3, Utf1, Tfap2c* were reported to maintain the pluripotency state (*32, 33*); *Dnmt3l, Dnmt3b* are DNA methylases, while *Tet1* is a demethylation enzyme (*32*); *Epcam* and *Cdh1* are mesenchymal–epithelial transition genes (*34*). In total, 60% (27 out of 44) pluripotency genes were significantly upregulated during reprogramming according to the mouse signature. On the other hand, no markers of pluripotency have been detected across significantly downregulated genes with the exception of 3 genes (*Ccnd1*, *Ccnd2* and *Cdkn1a*) known to be activated during the early stage of reprogramming and suppressed afterwards (*35*). Such enrichment of pluripotency-associated genes within the subset of upregulated, but not downregulated, genes indicates that our signature correctly characterizes the reprogramming process.

Finally, to compare reprogramming-associated gene expression changes across species, we examined the intersection of statistically significant genes (BH-adjusted p-value < 0.05) from the mouse and human cell reprogramming signatures (Fig. 1E). Fisher’s exact test showed significant co-regulation of genes during reprogramming in different species (p-value < 10e-10). In particular, 4 out of the top 5 upregulated mouse genes (*Parp1, Rcor2, Jarid2, Epcam*) were also significantly upregulated in the human signature. Similarly, 3 out of the top 5 downregulated mouse genes (*Zmpste24, Msrb3, Tbc1d8b*) were significantly downregulated in the human signatures. Therefore, although there are certain species-specific reprogramming features (*36*), this process appears to be highly similar in human and mouse cells at the level of gene expression. The obtained signatures allow investigating the interplay between molecular mechanisms of reprogramming and other traits, including aging and longevity.

### Reprogramming signatures are associated with biomarkers of longevity and aging

To explore the association between reprogramming, aging and longevity, we expanded our analysis with the gene expression signatures of mammalian aging and established lifespan-extending interventions identified previously (*27*). Aging signatures represent age-related gene expression changes in individual organs (liver, brain, muscle) of mice, rats and humans; common changes across different tissues within a certain species (mouse, rat, human), and a global signature characterizing common age-related changes across different tissues and species. Signatures of longevity interventions include biomarkers of individual lifespan-extending interventions (caloric restriction (CR), rapamycin, growth hormone (GH) deficiency), common biomarkers of interventions (Common) and genes, whose level of expression is correlated with mouse median and maximum lifespan (Median; Maximum) (*27*).

We observed significant positive correlations between several signatures of longevity interventions and reprogramming (mean rho = 0.11, p.adjusted < 0.05) (Suppl. Fig. S3). At the same time, aging-related changes demonstrated substantial negative correlation with both reprogramming and lifespan-extending interventions (mean rho = −0.13, p.adjusted < 0.05). As expected, the reprogramming signatures, including human-specific, mouse-specific and the combined signature, clustered together, pointing to the general similarity of this process across species. Aging and longevity signatures also formed separate clusters. Interestingly, reprogramming-associated changes clustered together with established lifespan-extending interventions, suggesting that in general reprogramming indeed recapitulates molecular mechanisms of longevity. Thus, clustering analysis of signatures agreed with the longevity and rejuvenation effects induced by reprogramming (Suppl. Fig. S3).

We next aggregated signatures within groups (Reprogramming, Aging, and Interventions) into combined meta-signatures to measure statistical significance of their co-regulation. There was a significant enrichment of co-regulated genes associated with reprogramming and longevity interventions (Fisher’s exact test, p-value = 0.00027, Fig. 2A, left panel), providing additional evidence of functional coherence of these two processes. Moreover, this co-regulation was preserved even after removal of all pluripotency genes or epithelial-mesenchymal transition genes from the analysis (not shown). This suggests that the longevity-associated effect of reprogramming may be uncoupled from pluripotency or the somatic identity program. Since reprogramming and aging signatures demonstrated significant negative correlations in our clustering analysis (Suppl. Fig. S3), we did not expect to find an enriched overlap of genes showing the same direction of expression dynamics between aging and reprogramming. Consistently, we observed a rather opposite, although not statistically significant, trend (Fisher’s exact test, p-value = 0.21, Fig. 2A, right panel).

**Fig. 2.**
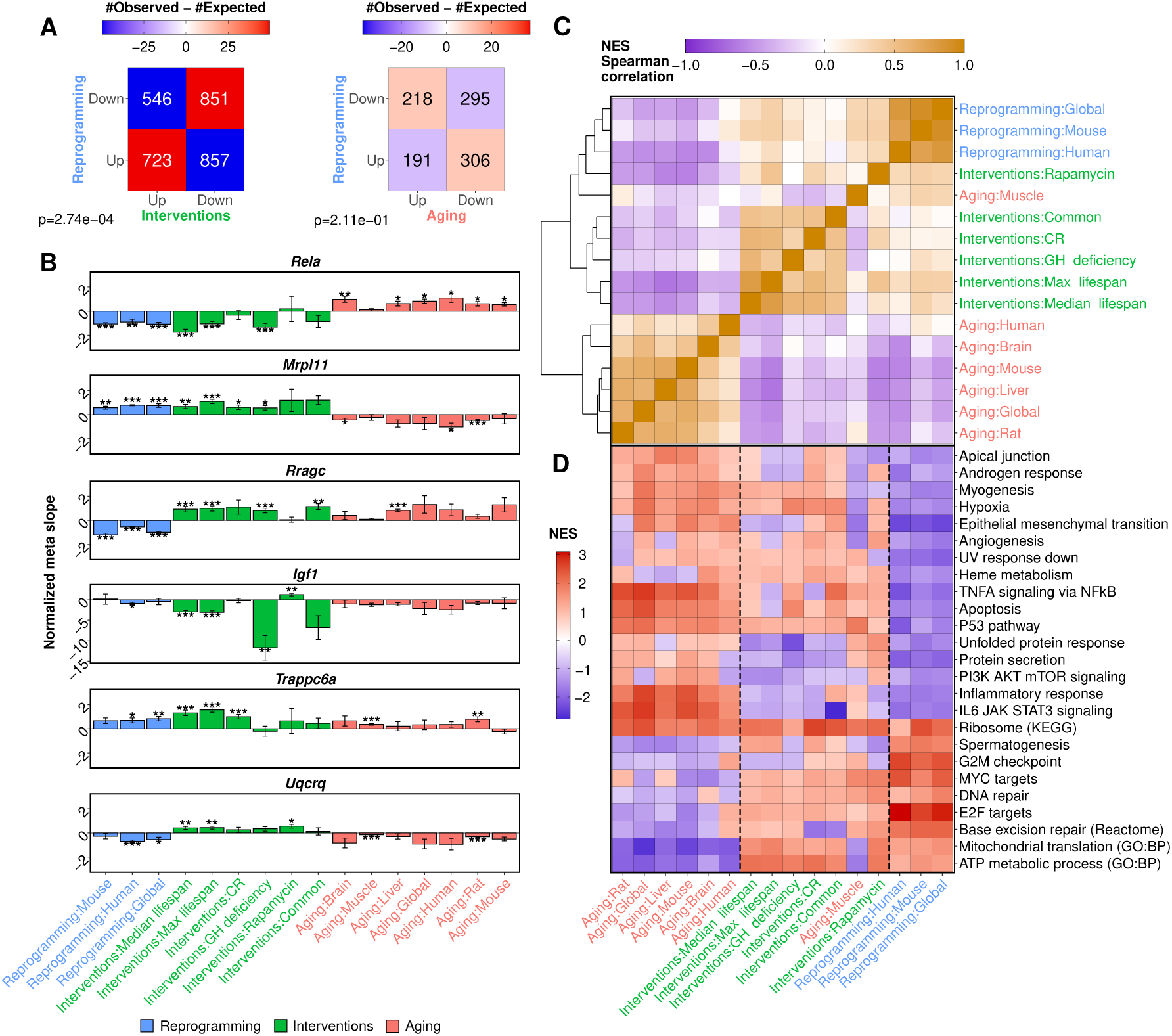
The interplay between reprogramming, aging and longevity signatures. **(A)** The overlap of upregulated and downregulated genes between groups of signatures. Numbers within cells show the observed numbers of overlapping genes, while color reflects the difference between observed and expected values. p-values are calculated using Fisher’s exact test. CR: caloric restriction, GH: growth hormone. **(B)** Barplots demonstrating the behavior of six particular genes across different signatures. Error bars represent standard errors of normalized meta-slopes. Annotation: * p.adjusted < 0.05; ** p.adjusted < 0.01; *** p.adjusted < 0.001. (C) Spearman correlation matrix of Normalized Enrichment Scores (NES) obtained using GSEA. **(D)** Functional terms across different signatures. Color represents NES values. Only the terms with at least one significant enrichment (adjusted p-value < 0.1) are shown. Dashed lines separate reprogramming and aging signature groups from the others.

We examined specific genes responsible for the discovered associations (Fig. 2B). For each group of signatures (Reprogramming, Aging and Interventions), we selected top genes with the lowest geometric mean of p-values. The first gene well-known for its association with aging and longevity, *Rela*, was downregulated upon reprogramming and in response to longevity interventions and upregulated during aging. *Rela* is a proto-oncogene, encoding a subunit of NF-*κ*B, and its human ortholog is known to influence age-related inflammation (*37*). The role of *Rela* downregulation during reprogramming is coupled with inhibition of NF-*κ*B pathway, which was reported to block a successful reprogramming in aged and progeria cells (*38*).

*Mrpl11* encoding the 39S subunit component of mitochondrial ribosome showed the opposite behavior, being positively regulated during reprogramming and by longevity interventions but downregulated with age. According to the GenAge database (*28*), deletion of this gene in *S. cerevisiae* decreases lifespan (*39*), suggesting that its level may affect longevity. However, the precise mechanistic role of this gene during aging and reprogramming remains unknown.

The other interesting example is *Rragc*, which has positive expression dynamics in the case of interventions and aging, but is downregulated during reprogramming. Rragc participates in the relocalization of mTORC1 to the lysosomes and its subsequent activation by the GTPase Rheb (*40, 41*). *Rragc* upregulation in longevity interventions and in the aging liver signature can be explained by the duality of Rag-GATOR pathway mechanism (*42*), depending on the source of amino acids. Downregulation of *Rragc* during reprogramming may be associated with transient mTOR pathway suppression influencing autophagy process (*43*). Interestingly, although expression of the gene coding for Insulin Like Growth Factor 1 (*Igf1*) was downregulated by the established longevity interventions, it wasn’t significantly perturbed during reprogramming.

Of particular interest are the genes showing the same direction of expression in all three signature groups: e.g. upregulated *Trappc6a* (encoding a trafficking protein particle complex that tethers transport vesicles to the *cis*-Golgi membrane) (*44*). Surprisingly, we found one gene, *Uqcrq*, with negative dynamics in aging and reprogramming, and positive in longevity interventions (Fig. 2B). This gene encodes a subunit of ubiquinol-cytochrome C reductase complex III, which is part of the mitochondrial respiratory chain (*44*).

### Functional enrichment analysis reveals processes associated with reprogramming-induced rejuvenation

To reveal functional processes associated with reprogramming, aging and longevity, we conducted gene set enrichment analysis (GSEA) of identified signatures (*45*). Similar to the individual meta-slopes (Suppl. Fig. S3), functional changes induced by reprogramming and established lifespan-extending interventions generally demonstrated a significant positive correlation with each other (mean rho = 0.23, adjusted p-value < 0.05) and were negatively associated with age-related changes (mean rho = −0.22, adj. p-value < 0.05) (Fig. 2C). Remarkably, normalized enrichment scores of functions were correlated even stronger than meta-slopes of individual genes.

Certain functional terms well characterized the identified reprogramming signature (Fig. 2D). For example, we observed downregulation of genes related to the Epithelial mesenchymal transition (EMT), the process which was shown to be suppressed during reprogramming (*23, 46*). Among the pathways downregulated by reprogramming but upregulated with age, we observed several terms related to inflammation: Inflammatory response, IL6/JAK/STAT3 signaling pathway, TNF*α* signaling via NF*κ*B (adjusted geometric mean p-value < 0.007 for each term and signature group). On the other hand, terms corresponding to mitochondrial function (Mitochondrial translation, ATP metabolic process) were upregulated in response to lifespan-extending interventions and reprogramming but downregulated with age (adjusted p-value < 0.03 for each term and signature group). This analysis pointed to the specific cellular processes associated with the longevity and rejuvenation effects of reprogramming. However, reprogramming did not appear to be a typical longevity intervention. In particular, it did not induce upregulation of the p53 pathway (adjusted p-value = 0.004), one of the common biomarkers of lifespan-extending interventions. Besides, it was also associated with downregulation of certain pathways upregulated by longevity interventions (*27*), including Heme metabolism, Hypoxia and Apoptosis (adjusted p-value < 0.008 for each term in reprogramming group).

### Clustering analysis of gene expression dynamics during reprogramming reveals specific trajectories of longevity-associated genes

To investigate specific dynamics of expression of longevity-associated genes during reprogramming, we performed a clustering analysis. First, we aggregated time series datasets of iPSC generation in mouse cells and calculated average trajectory for each gene significantly perturbed during reprogramming (see Methods). Next, we clustered genes by their trajectory using an agglomerative clustering approach. This approach resulted in 4 gene clusters selected using the Elbow criterion (Fig. 4A,B). Two major clusters (2 and 3) included genes that were almost monotonously up- or downregulated with time, respectively. Consistent with the data in Figure 1C, their expression followed a hyperbolic trajectory with a characteristic saturation at approximately 10th day of reprogramming. The expression of genes from two other clusters (1 and 4) followed U-shaped curve, starting from a transient up- or downregulation, respectively, and gradually returning back to the initial expression value afterwards. These genes reached their peak expression value after approximately 4-6 days of reprogramming.

**Fig. 4.**
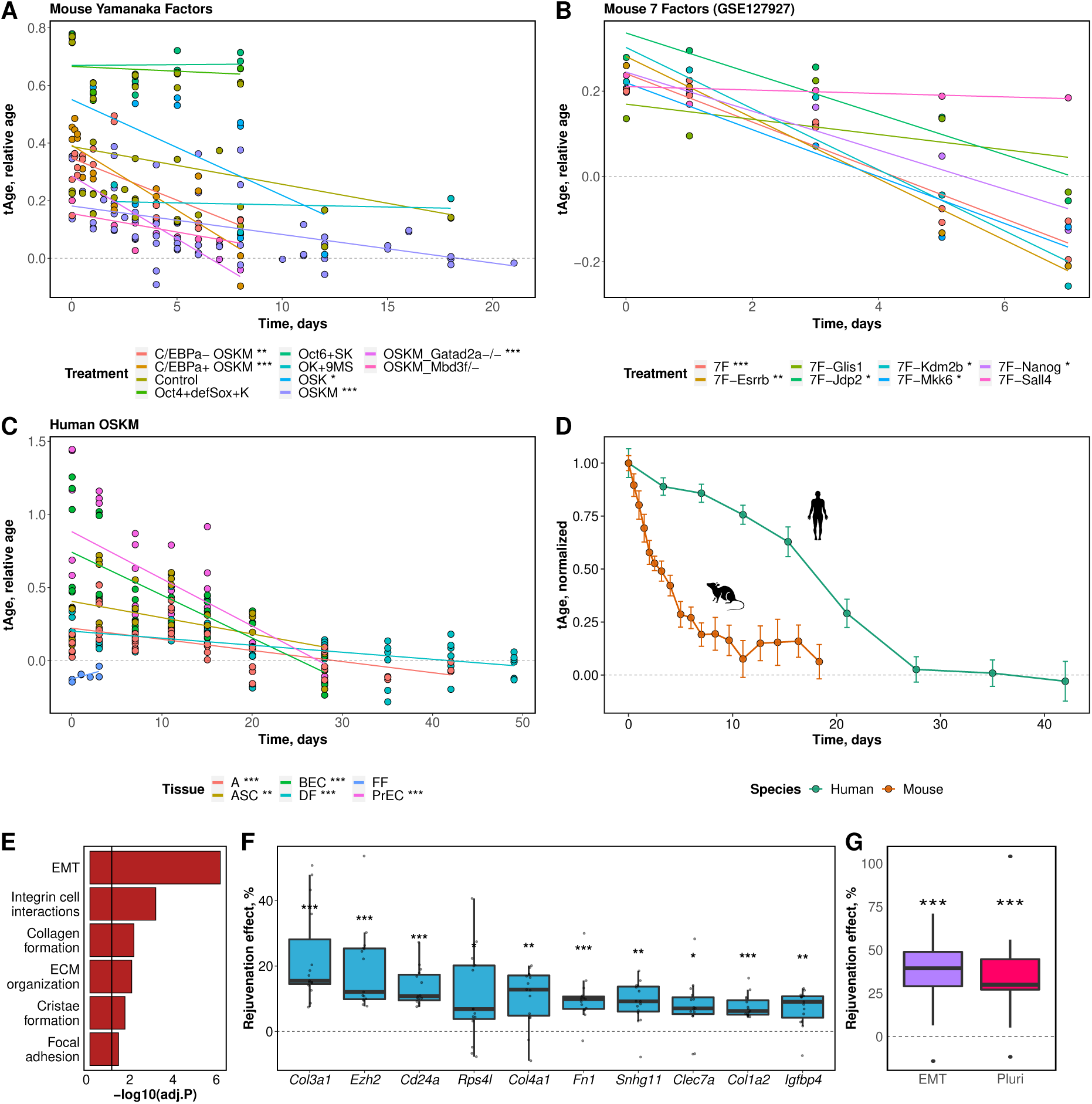
The evaluation of gene expression age during reprogramming using multi-tissue transcriptomic clocks. **(A)** Transcriptomic age (tAge) changes during reprogramming of mouse cells induced by Yamanaka factors. Lines represent the fitted linear model for the corresponding type of treatment. Stars indicate the significance level of the corresponding linear model slope. Relative age is defined as a chronological age divided by the maximum lifespan for a given species. **(B)** tAge changes during reprogramming of mouse cells induced by 7 factors from (*11*). All notations are the same as in panel A. **(C)** tAge changes during reprogramming of human cell lines. All notations are the same as in panel A. Each color represents a certain cell type. **(D)** Aggregated trajectories of rejuvenation induced by reprogramming for mouse and human cells. The curve is smoothed using moving average and normalized by the average tAge at the first point of time. Errorbars represent standard errors of the mean. **(E)** Functional enrichment analysis of genes from the mouse tClock model with significant effect on rejuvenation during reprogramming. The black line shows the significance threshold (adjusted p-value=0.05). Hallmark (epithelial-mesenchymal transition), KEGG (focal adhesion), Reactome (integrin cell surface interactions, extracellular matrix organization, collagen formation), and GO:BP (cristae formation) terms are presented on the barplot. **(F)** Distributions of rejuvenation effects of top genes associated with murine RIR across the datasets. The top 10 genes are sorted by their average contribution to rejuvenation (see Methods) according to the mouse tClock model. **(G)** The portion of RIR effect caused by the regulation of EMT (green) and pluripotency-associated (orange) genes. Each boxplot reflects the distribution across individual datasets. ASC: Adipose-derived stem cell, A: human astrocytes, BEC: bronchial epithelium cells, DF: dermal fibroblasts, FF: foreskin fibroblasts, PrEC: Prostate epithelium cells, EMT: Epithelial-Mesenchymal transition, Pluri: pluripotency-associated genes. * P<0.05, ** P<0.01, *** P<0.001

Then, each of the clusters was assessed for enrichment of longevity- and aging-associated genes obtained from the previously described signatures using Fisher exact test (Fig. 4C). Upregulated and downregulated signature genes were analyzed separately. The most significant association was observed between clusters 2-3 and signatures of longevity interventions, including biomarkers of CR and GH deficiency as well as genes associated with murine median and maximum lifespan signatures (adjusted p-value < 0.05). Remarkably, genes from both of these clusters were regulated by longevity interventions in the same direction. Thus, genes up- and downregulated during reprogramming (clusters 2 and 3) were enriched for genes up- and down-regulated in response to lifespan-extending interventions, respectively. Functional enrichment analysis of the clusters revealed that genes in 2 clusters upregulated during reprogramming included replication activating E2F target genes as well as genes involved in base excision repair and G2-M checkpoint (adjusted p-value < 0.0032). Cluster 3, downregulated during reprogramming, was enriched with genes related to EMT, Inflammatory response and Myogenesis (adjusted p-value < 0.001). Genes in clusters 1 and 4 were associated with several signatures including CR (co-regulation with 1) and aging in brain and in rats (opposite regulation with 1 and 4). Cluster 1 following U-shape behavior was enriched for genes involved in the TNF-alpha signaling pathway, Hypoxia, Protein secretion and Myogenesis (adjusted p-value < 0.003). Finally, cluster 4 demonstrating the opposite dynamics was functionally associated with G2-M Checkpoint, E2F targets, Myc targets, and mitochondrial translation (adjusted p-value < 0.003).

Thus, our cluster analysis of murine cell reprogramming suggests that genes monotonously changed during reprogramming show a significant co-regulated association with the biomarkers of lifespan extension. Interestingly, these genes were mostly perturbed during the first 6 days of reprogramming, suggesting that even transiently reprogrammed cells may acquire a longevity-associated transcriptomic phenotype, consistent with the experiments in vivo (*15, 16*).

### Transcriptomic clock reveals the rejuvenation effect of reprogramming

To estimate the systemic rejuvenation occurring during reprogramming, we utilized our recently developed mouse and human multi-tissue gene expression aging clocks (unpublished). These transcriptomic clocks (tClocks) were constructed based on more than 2,000 samples from 94 datasets across multiple tissues of mouse and human. We applied the clocks to predict the change of transcriptomic age (tAge) during reprogramming of mouse (Fig. 3A, B) and human cells (Fig. 3C). We also compared tClocks predictions with with epigenetic ages estimated using Horvath clock ((*21*)) utilizing the dataset with both DNA methylation and gene expression measured at once (*47*). We observed a significant positive correlation between the predictions (Suppl. Fig. S5), showing consistent behavior of clocks developed using different types of molecular data.

**Fig. 3.**
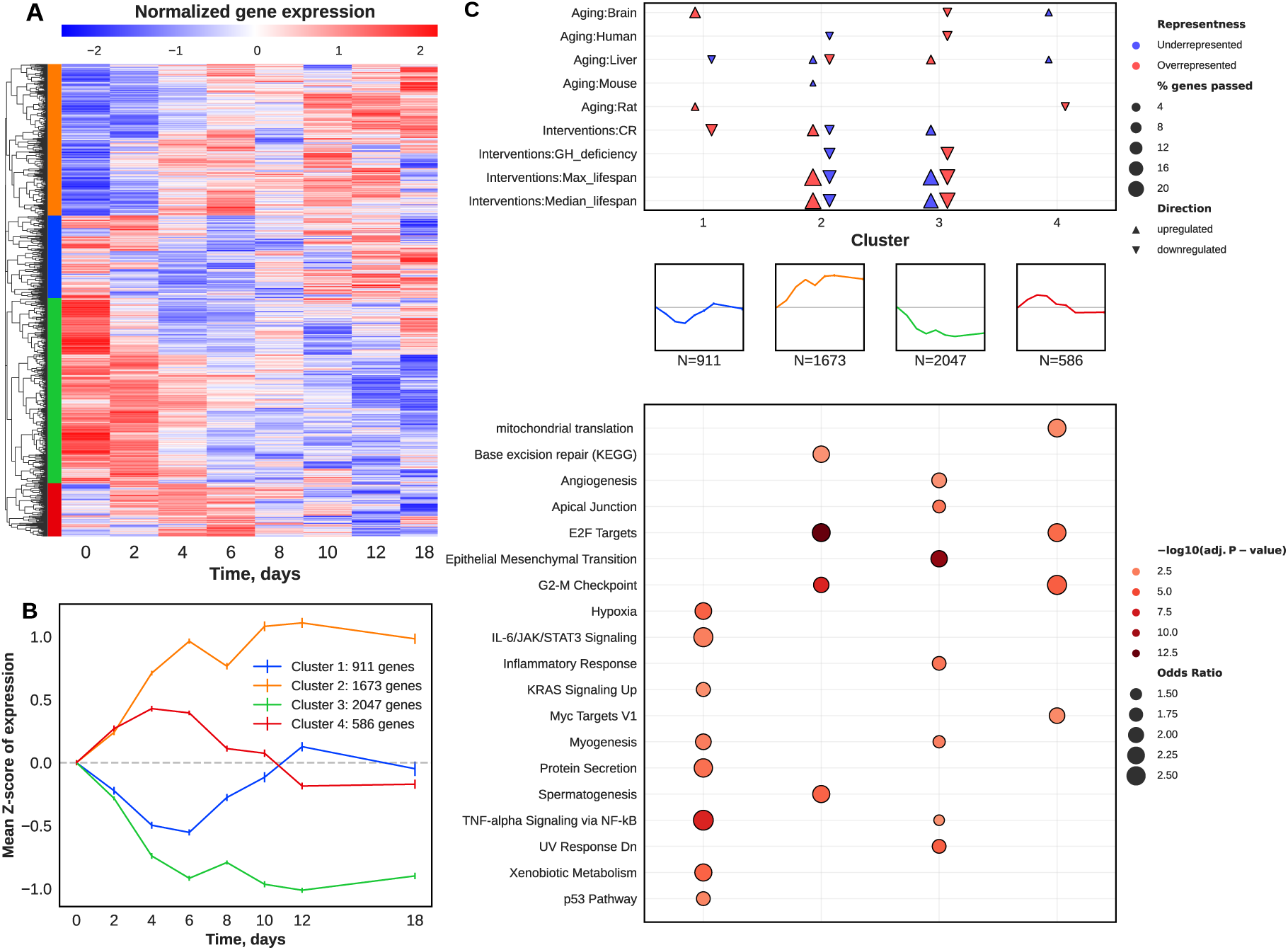
Clustering analysis of murine cell reprogramming-associated gene expression changes. **(A)** Normalized gene expression during reprogramming. Only genes with statistically significant non­ constant expression during reprogramming (ANOVA adjusted p-value < 0.05) are shown. Each row corresponds to a single gene. Colors represent four clusters chosen with the Elbow criterion. (B) Dy­ namics of cluster centroids during reprogramming. Error bars represent the standard error of the mean. The colors of clusters correspond to panel A. (C) Cluster enrichment analysis. Top panel: enrichment of clusters by genes associated with aging and longevity. Enrichment by up- and downregulated genes is reflected by the direction of triangles. The statistical significance of the overlap is assessed using Fisher’s exact test. Overrepresentation and underrepresentation are shown in red and blue, respectively. The absence of triangle reflects non-significant results (BH-adjusted p-value >= 0.05). Only signatures with significant enrichment in at least one cluster are shown. Triangle size represents the proportion of genes corresponding to a particular signature within a cluster. Middle panel: cluster centroids schematics (from B). Bottom panel: functional enrichment of clusters assessed with hypergeometric test. Only terms with significant enrichment in at least one cluster are shown. Color represents statistical significance of association, while size of the bubbles reflects odds ratio.

We found significant transcriptomic rejuvenation of murine cells during reprogramming induced by YF (Fig. 3A). Different variants of YF treatment, including OSKM and OSK, resulted in a significant decrease of tAge during reprogramming (p-value < 0.05), with the exception of OK+9MS (*48*) treatment and OSKM accompanied by *Mbd3f* knockout (*49*) (Suppl. Fig. S4A). Treatment of cells with the full set of 7 reprogramming factors also led to a decrease in tAge (p-value = 1.43e-05) (Fig. 3C, Fig. S4A). At the same time, one-by-one removal from the cocktail of these factors displayed diverse behavior. Specifically, removals of *Esrrb*, *Nanog*, *Mkk6*, *Kdm2b* (also known as *Jhdm1b*), *Jdp2* did not diminish the RIR effect, whereas removals of *Glis* and especially *Sall4* blocked the rejuvenation process. Interestingly, removal of *Sall4* at the same time resulted in a dramatic decrease in reprogramming efficiency (*11*). On the other hand, removal of *Esrrb* also led to a significant decrease in reprogramming efficiency but did not impair rejuvenation according to the transcriptomic clock, suggesting that the rejuvenation effect can be at least partly decoupled from the pluripotency state induction. Remarkably, the final tAge of reprogrammed cells subjected to YF and 7F was close to 0 for most datasets (average tAge = −0.0087 on day 19), which is consistent with the epigenetic data (*19*). It suggests that features of aging are reset during reprogramming both at the gene expression and DNA methylation levels.

To explore specific genes, whose expression change resulted in RIR, we measured the change in tAge after removing each individual gene from the mouse tClock model (see Methods), further referred to as a rejuvenation effect (RE) of a gene. We calculated RE for all genes with non-zero coefficients in the model (337 genes in total) across all datasets where significant rejuvenation was observed (adjusted p-value < 0.05). We identified 84 genes with the positive and significant (adjusted p-value < 0.05) rejuvenation effect. Enrichment analysis of this set of genes indicated a strong relation to Epithelial-Mesenchymal Transition (EMT) and processes involved in Extracellular matrix organization, including Collagen formation, Integrin cell surface interaction, and others (adjusted p-value < 0.05 for all presented terms) (Fig. 3E).

Next, we searched for genes contributing primarily to rejuvenation according to the mouse tClock model. Surprisingly, only one pluripotency-associated gene - *Ezh2* - was found among the top 10 predictors of RIR (Fig. 3F). However, Polycomb-group gene *Ezh2* (*50*) contributed 19% of the total rejuvenation effect on average across datasets (adjusted p-value=0.0004). Other genes in the top 10 were associated mostly with EMT (i.e., *Col3a1*, *Igfbp4*, *Postn*, *Fn1*). We further assessed the RE after removing all genes related to EMT or pluripotency from the model (Fig. 3G). We observed a significant reduction of the rejuvenation effect by 37% on average after removing EMT genes (p-value = 1.193e-05) and 35% on average after removing pluripotency-associated genes (p-value = 1.18e-04). It’s worth noting that EMT and pluripotency gene sets have no common genes. These results provide an estimate of the impact of pluripotency and EMT related genes on reprogramming-induced rejuvenation, suggesting that the major part of RIR is not explained by the perturbed expression of genes associated with somatic identity.

An analogous analysis of human cell reprogramming following OSKM treatment produced similar results (Fig. 3C). Using a human multi-tissue tClock, we observed a significant rejuvenation (adjusted p-value < 0.05) in almost all cell lines during reprogramming. The only exception was a dataset on foreskin fibroblasts containing very few data points. Interestingly, the rejuvenation effect of individual genes demonstrated high variance across human cell lines (data not shown), suggesting that the rejuvenation process during reprogramming may be achieved through regulation of various genes depending on the tissue. Consistently, human rejuvenating genes were not significantly enriched in any functional terms, supporting high heterogeneity of RIR across tissues.

To compare the rejuvenating trajectories of human and mouse cells subjected to OSKM treatment, we aggregated normalized tAge values across the datasets for each species and applied a moving average smoothing approach (Fig. 3D). We observed a rapid decrease of transcriptomic age of murine cells following the shape of exponential decay. On the other hand, rejuvenation of human cells followed a sigmoid curve. Although the transcriptomic age of cells from both species was close to zero at the end of reprogramming, the RIR of human cells required more time. Remarkably, this time difference was consistent with the duration of the reprogramming process, lasting, on average, for 14 and 30 days for mouse and human cell lines, respectively (*36*).

### Reprogramming signature uncovers new geroprotective interventions

To identify treatments that induce reprogramming-associated rejuvenation at the gene expression level, we selected genes displaying contrasting expression patterns according to aging and reprogramming signatures (Fig. 2A, right panel). We then used these genes as a query for the Connectivity MAP (CMAP) database (*51*) (see Methods). CMAP database contains gene expression profiles of human cells treated with different genetic or chemical interventions. CMAP connectivity analysis provides connectivity scores as a measure of similarity between a given gene set and transcription changes induced by perturbations from the database.

We selected top 20 perturbations showing the most significant positive or negative association with the reprogramming-associated rejuvenation signature for each type of perturbation: over-expression of a particular gene, treatment with a particular compound, knockdown of a gene via shRNA, and knockout of a gene via CRISPR-Cas9 system. To validate the rejuvenation effect of identified interventions, we applied the human and mouse aging tClocks to gene expression profiles of untreated and treated samples from the CMAP database separately for each available cell line. We then aggregated the obtained tAge values across cell lines using linear regression models (see Methods). In the end, we obtained two estimates of rejuvenation effects for each treatment, including aggregated connectivity scores from the CMAP analysis and aggregated tAge values from aging clocks (Fig. 5A).

**Fig. 5.**
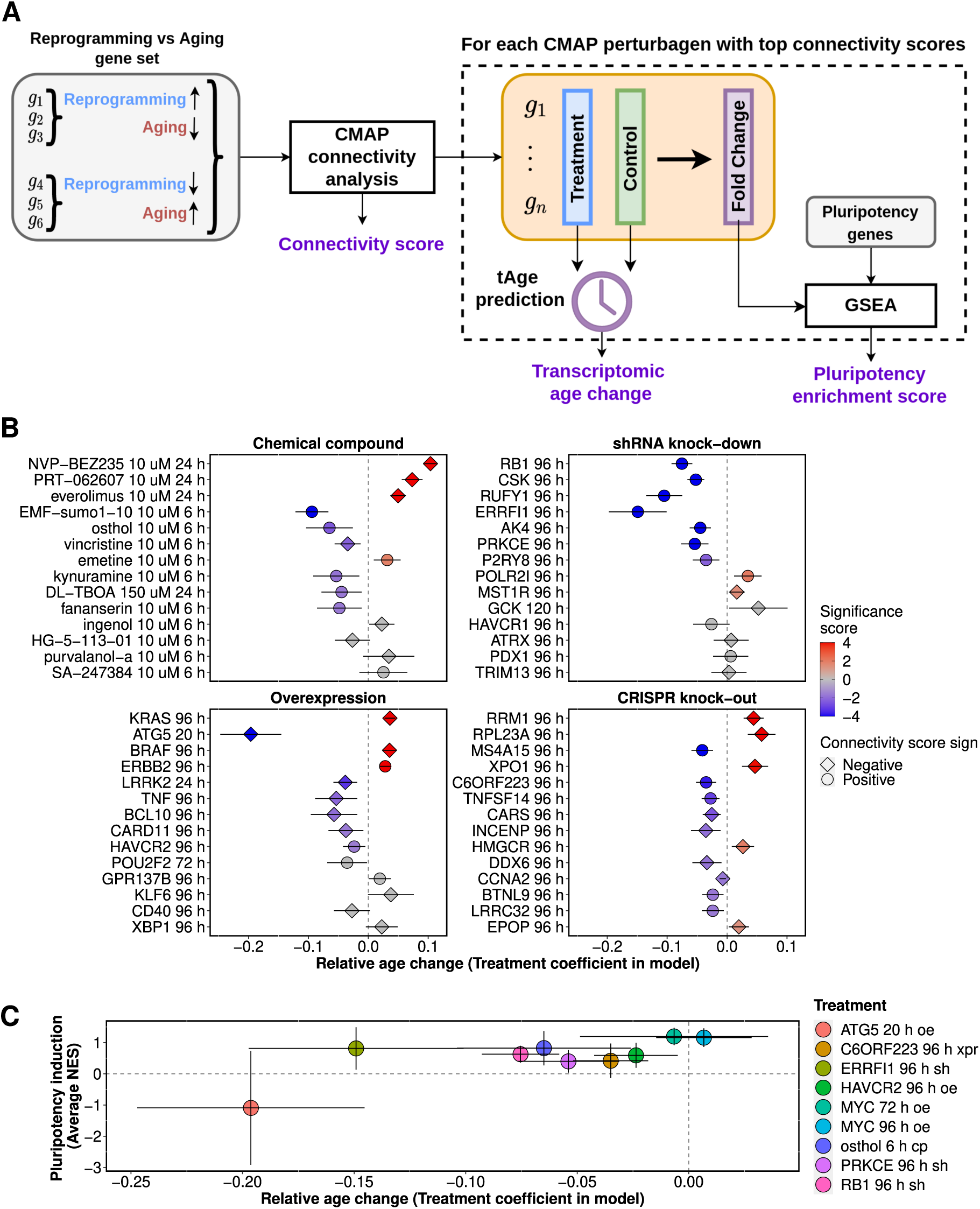
Identification of rejuvenation-associated interventions using CMAP. **(A)** Schematic illustration of CMAP analysis workflow. **(B)** Relative age change for different types of interventions. Age change corresponds to the *Treatment* coefficient from the aggregation model (see Methods). Whiskers correspond to the 95% confidence interval. The significance score is computed as −Log10(P-value) * Sign(Treatment coefficient). Grey points indicate insignificant coefficients. Circles correspond to coinciding directions of signature connectivity and tClock analysis, while other symbols correspond to opposite directions. **(C)** Rejuvenation- and pluripotency-inducing effects of selected interventions. Rejuvenation effect was assessed using tClock while pluripotency effect was determined with GSEA. Whiskers correspond to the 95% confidence interval. oe: overexpression, NES: Normalized Enrichment Score, h: hours, sh: gene knockdown with short hairpin RNA, xpr: gene knockout with CRISPR.

Among interventions that demonstrate significant rejuvenation effects based on the connectivity score as well as both mouse and human tClocks we observed overexpression of *HAVCR2* (*TIM3*), which encodes a cell surface receptor implicated in modulating innate and adaptive immune response (adjusted p-value = 0.028). Interestingly, overexpression of *TIM3* was shown to alleviate inflammation in human patients with thyroid-associated ophthalmopathy via suppressing the Akt/NF-*κ*B signaling pathway (*52*). At the same time, *Tim3* overexpression resulted in deterioration of neuroinflammatory and neurocyte apoptosis in a rat subarachnoid hemorrhage model (*53*). Together, these observations suggest that *TIM3* plays a significant role in regulation of age-associated inflammatory processes, and its overexpression can be considered as a treatment against inflammaging.

Knockdown of *ERRFI1* (*MIG-6*) was also found to decrease the cellular transcriptomic age according to both our clocks (adjusted p-value = 1.24e-8). Expression of *ERRFI1*, which encodes a negative regulator of EGFR signaling, is upregulated during the cell growth (*54*). Interestingly, overexpression of *MIG-6* was shown to be sufficient to trigger premature cellular senescence (*55*). In contrast, knockdown of *MIG-6* delayed the initiation of Ras-induced cellular senescence (*56*), supporting our conclusion derived from tClock.

Among top interventions inducing a significant rejuvenation effect across different cell lines according to the human clock, we identified knockdown of *PRKCE* (adj. p-value = 1.7e-5), knockout of *C6ORF223* (adj. p-value = 1.9e-4) as well as a treatment with a chemical compound osthol applied at 10 uM dose for 6 hours (adj. p-value = 0.0038). Interestingly, osthol has been shown to demonstrate anti-inflammatory effects by blocking the activation of NF-*κ*B and MAPK/p38 pathways (*57*). In addition, *osthol* prevents accumulation of advanced glycation end products (AGE) via the induction of *Klotho* expression (*58*). The rejuvenation effect of *PRKCE* knockdown and *C6ORF223* knockout was also supported by experimental data. Thus, inhibition of *PKC* signaling was shown to maintain self-renewal and pluripotency of rat embryonic stem cells (*59*), while *C6ORF223* accumulation was associated with age-related macular degeneration (*60*) and was correlated in expression levels with a well-known human aging-related gene *VEGF* (*61, 62*). Remarkably, one of the top interventions predicted by our model was over-expression of *ATG5* showing a strong rejuvenation effect by human clocks (adjusted p-value = 1.75e-12). *ATG5* gene product is involved in autophagy, mitochondrial quality control, regulation of the innate immune response and other cell processes. In fact, *ATG5* overexpression was shown to increase lifespan of healthy mice by enhancing autophagy (*63*).

Finally, to investigate whether the treatments described above induce expression of pluripotency-associated genes, we performed GSEA analysis testing if genes differentially expressed in response to interventions are enriched for pluripotency genes from (*64*) (see Methods). After obtaining NES scores for each cell line, we aggregated them using average and applied t-test to assess significance of the aggregated score. As a positive control, we performed a similar analysis in 2 models of overexpression of *MYC*, known to partially induce the pluripotency program in cells (*9, 22*). The latter treatment showed a significant upregulation of pluripotency genes (p-value < 8.94e-5), while it did not result in a significant reduction of cellular tAge. On the other hand, *ATG5* overexpression induced a rejuvenation effect according to tClock without activation of pluripotency genes. In fact, it even slightly suppressed pluripotency program, though insignificantly (p-value = 0.083). Similar significant rejuvenation combined wifth neutral effect on pluripotency (p-value = 0.128) was produced by knockout of lncRNA *C6ORF223*. These examples confirm that rejuvenation and loss of somatic identity associated with reprogramming can be decoupled, and interventions separately affecting each of these processes may be developed.

## Discussion

Reprogramming-induced rejuvenation is a fundamental concept denoting a family of cell reprogramming approaches focused on their capacity for rejuvenation (*65*). These approaches gained much attention in recent years as they have the potential for radical interference into aging and longevity (*2–7*). Therefore, it is essential to understand precisely which processes during reprogramming lead to rejuvenation and how they can be decoupled from the loss of somatic identity. In this study, we investigated these processes and provided a systemic view on rejuvenation during reprogramming by analyzing signatures identified from multiple time-course reprogramming datasets and revealing their interplay with biomarkers of aging and lifespan extension (Fig. 6A).

**Fig. 6.**
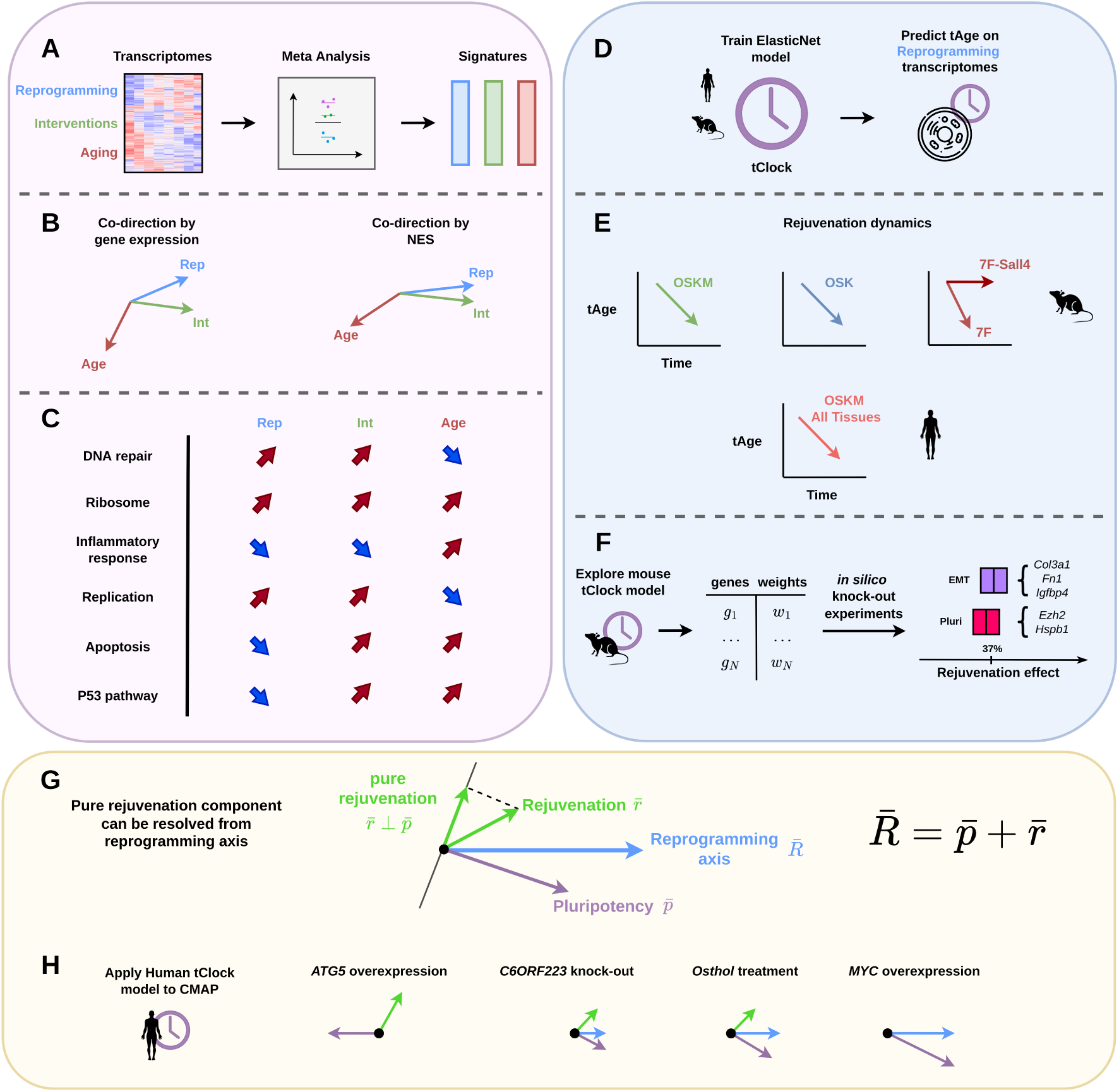
Summary of the work. **(A)** Gene expression signatures of cellular reprogramming in mouse and human were constructed via aggregation of multiple datasets by meta-analysis technique. **(B)** Reprogramming signature was positively correlated with biomarkers of longevity interventions and negatively correlated with signatures of aging (meta-slopes correlation). The strength of these associations was amplified at the functional level (NES correlation). **(C)** Functional behavior of signatures in selected ontological terms. Red and blue arrows denote positive (upregulation) and negative (downregulation) NES scores, respectively. **(D)** tClock models trained on the mouse or human aging gene expression datasets were used to predict the tAge of reprogramming cells across time. **(E)** OSK, OSKM Yamanaka factors, and 7F factors consistently decrease tAge of murine and human cells during reprogramming. However, removal of certain factors, such as Sall4, may result in abrogation of both reprogramming and rejuvenation. **(F)** Genes contributing to the rejuvenation effect of reprogramming were identified. Pluripotency and EMT-associated genes are responsible for approximately 35-37% of RIR. **(G)** Reprogramming can be considered as a vector being a sum of two components: one moving cell toward pluripotency and the second moving cell to a rejuvenated phenotype. Investigation of gene expression signatures allows to decouple these processes. **(H)** New interventions affecting one of the reprogramming-induced components can be discovered using instruments provided in this work. Rep: Reprogramming; Int: Lifespan-extending interventions; Pluri: Pluripotency; EMT: Epithelial-mesenchymal transition.

We were able to construct robust reprogramming signatures and show that: (i) mouse and human signatures are well correlated with each other and share a significant number of genes regulated in the same direction (Fig. 1E); and (ii) reprogramming signatures are positively correlated with longevity interventions and negatively correlated with various aging signatures. In addition, we discovered co-regulation of particular genes in response to reprogramming and established lifespan-extending interventions including downregulation of *Rela* and upregulation of *Mrpl11* previously shown to be significant biomarkers of murine longevity (*28,37,38*). The associations between three groups of signatures - reprogramming, interventions, and aging - persist and are even amplified at the level of functional enrichment (Fig. 2C, 6B). Most conspicuous functions (Fig. 6C) demonstrate that reprogramming may act as a longevity intervention but not in all aspects. Of note, these results are generally consistent with those obtained from single-cell analysis (*22*). Namely, we observed that the reprogramming suppressed genes were associated with inflammatory response. On the other hand, upregulation of fatty acid metabolism observed after transient reprogramming of mouse mesenchymal stem cells was not prominent in our signatures. Multiple studies have previously demonstrated that the DNA methylation age (mAge) decreases during the reprogramming process (*15, 18, 19, 66*). However, only a few studies (*20*) attempted to reproduce these results at the transcriptome level using single-tissue clocks (see (*67*) for details). To fill this gap, we utilized mouse and human clocks trained on multiple tissues to predict transcriptomic age (tAge) of cells during the whole reprogramming process (Fig. 6D). As expected, we observed a systematic decrease of tAge for the majority of reprogramming datasets for both mouse and human cell lines (Fig. 6E, 3C,D). Notably, some treatments that failed to result in successful reprogramming during the original experiment (e.g. 7F-Sall4, (*11*)), did not lead to the decrease of tAge with time. On the other hand, some of the treatments that didn’t lead to the gain of pluripotency significantly decreased transcriptomic age of somatic cells (e.g. 7F-Esrrb, (*11*)). This is consistent with results of (*22*) reporting that induction of only SK factors decreases aging score without loss of mesenchymal identity. Such results support the possibility of decoupling reprogramming-induced rejuvenation from the changes involved in the loss of somatic identity.

Next, we explored genes responsible for RIR by conducting *in silico* knockout experiments. We identified several genes that contributed the most to the rejuvenation process. Among the top 10 genes, there was only one pluripotency-associated gene - *Ezh2* (Fig. 3F). We also observed several genes associated with EMT, e.g., *Col3a1*, *Igfbp4*, *Postn*, *Fn1*. In total, 37% of RIR, on average, was explained by the EMT genes, and 35% of the RIR was affected by the pluripotency-associated genes. Therefore, the tClock model suggests that although a part of the RIR is achieved through the deregulation of genes involved in the maintenance of somatic identity, a significant portion of it is orthogonal to this process. Genes responsible for that effect represent perspective biomarkers allowing to search for new geroprotectors.

To discover such interventions, we conducted CMAP (*51*) connectivity analysis and revealed treatments that produced rejuvenation-associated gene expression changes similar to reprogramming. We validated our hits using human transcriptomic clocks and revealed several interventions with a potential rejuvenation effect. Consistently, some of them, including *ATG5* overexpression, *C6ORF223* knockout and osthol treatment, have been previously shown to have a positive effect on lifespan (*57, 58, 60, 63*). In addition, we tested these and other rejuvenating interventions *in silico* for their ability to induce pluripotency program (5C) and observed that *ATG5* overexpression and *C6ORF223* knockout did not significantly affect the expression of these genes across multiple cell lines (Fig. 5H). Therefore, according to our data, these treatments appear to produce RIR without affecting somatic cell identity. Interestingly, *MYC* over-expression demonstrated the opposite effect, producing no significant rejuvenation effect but inducing the pluripotency program, in agreement with its role as one of Yamanaka’s factors but interfering with results of (*22*) where induction of this factor showed little loss of mesenchymal identity but also small decrease in aging score.

Taken together, these results indicate that the reprogramming process contains a rejuvenation component that can be expressed in the gene or function dynamics. Recent *in vivo* reprogramming demonstrated no systemic rejuvenation of all murine tissues with the exception of skin and kidney tissues (*18*). The authors hypothesize that this is due to some tissues being more susceptible to OSKM reprogramming than others. It can even be assumed that the OSKM set of factors may not be suitable for *in vivo* reprogramming. Moreover, the fact that this set of factors is known to be oncogenic forces researchers to develop complex treatment protocols. This complexity can be avoided if the oncogenic aspect is completely excluded, which is proposed in the RIR concept. Today, several of the possible ways to solve this problem include the use of OSK reprogramming (*15*), reprogramming until the maturation phase achieved (*20*) or even chemical reprogramming (*12*). However, understanding the mechanisms of rejuvenation achieved during reprogramming may provide us with better solutions. Future rigorous studies should reveal which gene networks are responsible for the regulation of the RIR process. Analysis of epigenetic aspects of RIR, such as methylation or histone modifications accompanying expression dynamics, may be a future direction. The ultimate solution would be to construct a dynamic mathematical model of RIR to predict not only a subset of transcription factors (or small molecule compounds) but also other characteristics necessary for successful treatment.

We propose a geometric metaphor to better represent the essence of rejuvenation during cell reprogramming (Fig. 6G). We represent the reprogramming process as a vector 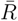 in the space of transcriptomic signatures. We assume that this vector can be decomposed into two non-orthogonal components: rejuvenation 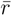 and pluripotency 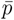 (here we mean the cumulative signature towards pluripotency). Their sum gives the original reprogramming vector 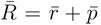. It follows from our analysis that pluripotency may proceed without rejuvenation (Suppl. Fig. 3A), and rejuvenation can occur without successfully achieved pluripotency (exemplified by the 7F-*Essrb* treatment Fig. 3B). It means that rejuvenation and pluripotency have co-directed components (projections onto the reprogramming vector) and also have orthogonal components (e.g., projection of rejuvenation vector onto the axis orthogonal to pluripotency). We argue that for successful rejuvenation without the risk of pluripotency-induced tumorigenesis, we need to identify the signature of “pure rejuvenation” 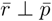, i.e., a set of genes with corresponding levels of expression that causes cell rejuvenation without notable shift towards pluripotency. The results obtained in this study using transcriptomic clock suggest that such genes include *Col3a1*, *Fn1*, *Cd24a*, while *Ezh2* is an example of a gene contributing to rejuvenation but being also a marker of pluripotency. Using a multitool of transcriptomic signature analysis, we made a step towards decomposing rejuvenation and pluripotency vectors that may lead to the safe and efficient reprogramming-induced rejuvenation.

## Methods

### Data collection

We collected publicly available cell reprogramming datasets containing more than three time points across the reprogramming process (Table S1). ESC and iPSC states were excluded since they were not corresponding to any particular time point of reprogramming. We used only preprocessed data (e.g., read counts) provided in the GEO database by datasets’ contributors.

### Data preprocessing

To aggregate multiple datasets into a joint signature, we utilized an approach as in our earlier work (*27*). It consists of several steps (Fig. 1A). First, each dataset was normalized using a conventional normalization technique appropriate for the given data type. RLE normalization followed by log transformation was applied for RNA-seq data. Log transformation of intensities followed by scaling and quantile normalization was used for microarray data. Second, for each gene changing its expression value with time, a linear regression model was constructed using the *limma* package (*68*). Third, slope coefficients, their standard errors and related statistics were extracted from models and used to represent corresponding gene regulation (positive or negative). Thus, a positive or negative slope corresponds to an increasing or decreasing expression of a particular gene with time in a given dataset, respectively. Finally, slope values from different datasets were aggregated using the mixed-effects model constructed by the *metafor* package (*69*), with GEO ID introduced as a random term. For every gene, this model produced a meta-slope, being a weighted average of slopes across all analyzed datasets. Corresponding p-values were adjusted for multiple comparisons using Benjamini-Hochberg (BH) approach (*70*). Genes with adjusted p-value < 0.05 were considered significant and included in the final reprogramming signature.

### Selection of datasets for the aggregated signature

The critical step for constructing a correct aggregated signature is filtering out non-concordant datasets. We used the Spearman correlation of dataset slopes as a concordance measure and calculated pairwise correlation coefficients between all 29 mouse and 12 human datasets using the union of top 350 genes in each dataset ranked by the correlation p-value of slopes. The threshold of 350 genes was identified to be optimal for noise removal since it maximized the number of significant pairwise correlations (Benjamini-Hochberg adjusted p-value < 0.05 and absolute *ρ* > 0.1). Finally, we used the agglomerative clustering approach based on the Euclidean distance with complete linkage to extract the largest cluster among all 29 mouse and 12 human datasets (Suppl. Fig. S1). As a result, 19 out of 29 mouse datasets and 11 out of 12 human datasets passed the selection criteria (formed a dominant cluster, see also supplementary figure S1 and Methods).

### Signature construction

Prior to signature construction, we normalized slope coefficients from different datasets based on the following algorithm. First, Spearman correlation of reprogramming-related gene expression changes was calculated for each pair of datasets. For that, we obtained the top 350 statistically significant reprogramming-associated genes ranked by the correlation p-values in each dataset and then formed a union of two such gene lists within a pair of datasets. Then, multiple Deming regression was calculated simultaneously for each pair of datasets with significant correlations using the union of top 350 genes. During this step, the cumulative squared loss across all significantly correlated pairs of datasets within a certain signature was minimized using the L-BFGS-B method in the R function *optim*. Normalization coefficients were allowed to vary between 0.01 and 100. To establish the global minimum of the error function, multiple Deming regression was calculated 10 times with random initial sets of normalization coefficients, and final coefficients were chosen from the run with the smallest cumulative regression error. Among these 10 runs, the error minimum was the same for most runs, indicating that the global minimum was achieved for each signature.

Then we used the *rma.mv* function in the *metafor* package (*69*) to construct intercept-only multilevel mixed-effects model with nested random effects (*71*). As a response variable, we used Deming-normalized slopes derived for each dataset. Since datasets originated from diverse sources, we had to account for their heterogeneity across different experiments (i.e., different GSE IDs) and within the same experiment (i.e., the same GSE ID), implying the multilevel embedded structure of the model. Fixed effects were not considered within this model. The final model can be described with the following formula:

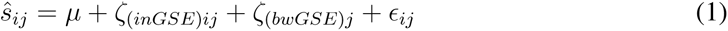

where 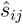 is an estimate of the true effect size *s_ij_*; *µ* is an actual mean of the slopes’ distribution; term *ij* denotes that some effect size *i* is nested in cluster *j*; *ζ*_(*inGSE*)*ij*_ is random term corresponding to a within-GSE-ID heterogeneity; *ζ*_(*bwGSE*)*j*_ is a random term corresponding to a between-GSE-ID heterogeneity; *ɛ_ij_* is a sampling error of individual datasets, which can be estimated from a standard error of a corresponding slope. We applied this model for construction of mouse, human, and combined signatures. Following the principle mentioned in the previous paragraph, we selected 19 datasets for mouse signature, 11 datasets for human signature, and 30 datasets (19 mouse + 11 human datasets) for the combined signature.

### Aggregation of p-values across the signatures

The aggregation of p-values within each group of signatures was conducted using the harmonic mean (*72*). Subsequently, we adjusted aggregated p-values using BH method. We selected significant genes (adjusted p-values < 0.05) to investigate the overlap of genes between reprogramming signature and signatures of aging and lifespan-extending interventions. The statistical significance of the overlap was assessed with Fisher’s exact test.

### Clustering analysis

To cluster genes by their expression dynamics, we first scaled all gene expression values, transforming them into z-scores. Next, we grouped observations into 2-day periods and applied one-way ANOVA considering 2-day intervals as a factor variable, testing a null hypothesis that average expression is equal over all intervals. Genes with the BH adjusted p-value < 0.05 were considered to demonstrate significant expression change over time. We excluded genes with constant expression and clustered the remaining genes using agglomerative approach with correlation distance metric and complete linkage, considering time intervals as features. The scikit-learn Python package (*73*) was used for this analysis.

### Prediction of transcriptomic age

To investigate the dynamics of gene expression biomarkers of aging during cellular reprogramming, we utilized multi-tissue transcriptomic mouse and human clocks based on signatures of aging across different tissues identified as explained in (*27*). The applied clocks were based on elastic net linear models that were designed to predict relative chronological age calculated as a real age divided by the maximum lifespan for a given species (48 months and 122 years for mouse and human, respectively). The missing values were omitted with the precalculated average values from the clock. Using the mouse and human clocks, we then calculated the transcriptomic age (tAge) for each mouse and human sample, respectively. Change of the tAge with time during reprogramming within each dataset was assessed using linear regression model. The slope of the tAge change with time was considered significant if the corresponding BH adjusted p-value < 0.05. For normalization of the tAge values across several OSKM-based reprogramming datasets, relative tAge values were divided by the average tAge value of the first time point within each dataset. After that, aggregated tAge trajectories for human and mouse data were smoothed using 3-day period moving average. Standard errors were calculated for smoothed tAge values in each time point.

### Estimation of rejuvenation effect of a gene

To estimate the rejuvenation effect of a specific gene in a particular dataset, the following pipeline was carried out: 1) *in silico* knockout was performed by making the expression of this gene equal to 0 for all of the samples; 2) tClock was used to calculate tAge for all samples in the given dataset before and after the “knockout”; 3) linear model was fitted to predict time-dependent tAge trajectory before and after “knockouts”; 4) the maximum difference between tAge estimates obtained from the linear model before and after “knockouts” was calculated; 5) the difference was normalized to the total rejuvenation effect in the dataset (the difference between the tAge value at the first time point and the tAge value at the final day of reprogramming). Thus, the result of this procedure demonstrates how the removal of certain gene affects the magnitude of tAge decrease during reprogramming, corresponding to its rejuvenating effect. The same approach was used to calculate the rejuvenation effect after “knocking out” the whole gene set (e.g., EMT or pluripotency-associated genes).

### Aggregated analysis of rejuvenation-inducing interventions based on CMAP

To identify treatments mimicking RIR at the gene expression level, we used CMAP query API (*51*). As a query, genes upregulated in combined reprogramming signature but downregulated in combined aging signature (’Up’ subset), and genes downregulated in combined reprogramming signature but upregulated in combined aging signature (’Down’ subset) were used. We will refer to these gene subsets as the RIR gene set.

The result of a CMAP query is essentially a list of perturbagens ordered by the score of association between differentially-expressed gene set and the query gene set. A positive score indicates a similarity between the query and effect of the given perturbagen applied to the certain cellular line, while a negative score indicates that these two signatures are the opposite to each other (i.e., genes that are increased by treatment with the perturbagen are decreased in the query, and vice versa). The magnitude of the score corresponds to the magnitude of similarity or dissimilarity between the treatment and query. Therefore, top and bottom hits in these lists represent interventions that have the strongest positive and negative associations with the query, respectively. These treatments appear to be of the highest interest for the subsequent investigation.

At the next step, we aggregated connectivity scores for each intervention with the same dosage and treatment time across different cell lines using simple averaging of connectivity scores. The statistical significance of the positive or negative association of the intervention across cell lines was assessed using t-test with the null hypothesis that the mean of connectivity scores across cell types is equal to zero. We then selected the top 20 positive and top 20 negative aggregated interventions from each of four intervention types (gene overexpression, chemical compound treatment, gene knockdown with shRNA, and gene knockout with CRISPR) for further analysis.

We downloaded gene expression data for the selected interventions and further applied transcriptomic clocks to the gene expression profiles induced by these treatments as well as control samples. Specifically, we obtained quantile normalized data from the CMAP level 3 data pre-processing step. We downloaded treatment data and corresponding control data for a given unique intervention-dosage-duration group indicator. After an additional data normalization procedure (see “Prediction of transcriptomic age” section for details), we applied the mouse and human transcriptomic clocks to the gene expression vectors. The obtained relative age values were aggregated with a linear model of the following form:

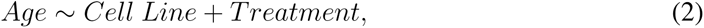

where *Age* is the relative tAge value, *C ell Line* is the name of corresponding cell line from CMAP (factor variable), and *T reatment* is the binary variable, which indicates whether the given relative tAge value is from the control or treatment subset. We fitted this model using *statsmodels* python package (*74*). The resulting coefficient of the *T reatment* variable can be interpreted as an average change in the relative tAge in response to a given intervention across cell lines, while its p-value reflects the statistical significance of this change. Thus, negative *T reatment* coefficient corresponds to “rejuvenation” effect while positive reflects “aging” effect. We paid particular attention to interventions with negative significant coefficient of the treatment variable coupled with the positive aggregated connectivity scores. Such interventions result in the gene expression response similar to reprogramming and opposite to aging and, at the same time, contribute to rejuvenation according to the transcriptomic clock, being of a particular interest.

Among the identified interventions, we searched for those not inducing the expression of pluripotency-related genes. First, we obtained differential gene expression data from the CMAP level 5 data preprocessing step for each of our top hits. Then, we performed gene set enrichment analysis (GSEA) using *fGSEA* package (*75*) testing if the gene expression response induced by a certain treatment is enriched for the set of pluripotency-associated genes obtained from (*64*). The calculated Normalized Enrichment Scores (NES) were then aggregated using simple averaging. The statistical significance of enrichment across cell lines was assessed using t-test with the null hypothesis that the mean of NES across cell types is equal to zero.

## Supporting information

Datasets used in the study

List of mouse and human pluripotency associated genes

Top 1000 genes from an identified reprogramming signature

## Acknowledgements

We thank Ruslan Gumerov for help with data collection. A.T. and S.E.D. were members of Interdisciplinary Scientific and Educational School of Moscow University “Molecular Technologies of the Living Systems and Synthetic Biology”.

## Author Contributions

A.T. and S.E.D. conceived and designed this research; D.K. performed research and data analysis; D.K., E.E.K, V.N.G., S.E.D. and A.T were involved in discussion and interpretation; D.K. and A.T. wrote the manuscript with contributions from all other authors.

## Data and materials availability

Additional data and materials are available online.

## Supplementary Materials

**Table S1. (separate file)** Datasets used in this study.

**Table S2. (separate file)** Table of pluripotency-associated genes for mouse and human.

**Table S3. (separate file)** Top 1000 genes from the mouse reprogramming signature sorted by statistical significance.

